# Central *in vivo* mechanisms by which *C. difficile’s* proline reductase drives efficient metabolism, growth, and toxin production

**DOI:** 10.1101/2023.05.19.541423

**Authors:** Laura M. Cersosimo, Madeline Graham, Auriane Monestier, Aidan Pavao, Jay N. Worley, Johann Peltier, Bruno Dupuy, Lynn Bry

## Abstract

*Clostridioides difficile* (CD) is a sporulating and toxin-producing nosocomial pathogen that opportunistically infects the gut, particularly in patients with depleted microbiota after antibiotic exposure. Metabolically, CD rapidly generates energy and substrates for growth from Stickland fermentations of amino acids, with proline being a preferred reductive substrate. To investigate the *in vivo* effects of reductive proline metabolism on *C*. *difficile’s* virulence in an enriched gut nutrient environment, we evaluated wild-type and isogenic *ΔprdB* strains of ATCC43255 on pathogen behaviors and host outcomes in highly susceptible gnotobiotic mice. Mice infected with the *ΔprdB* mutant demonstrated extended survival via delayed colonization, growth and toxin production but ultimately succumbed to disease. *In vivo* transcriptomic analyses demonstrated how the absence of proline reductase activity more broadly disrupted the pathogen’s metabolism including failure to recruit oxidative Stickland pathways, ornithine transformations to alanine, and additional pathways generating growth-promoting substrates, contributing to delayed growth, sporulation, and toxin production. Our findings illustrate the central role for proline reductase metabolism to support early stages of *C. difficile* colonization and subsequent impact on the pathogen’s ability to rapidly expand and cause disease.

## Introduction

*Clostridioides difficile* is a toxigenic, spore-forming, and anaerobic bacillus that can asymptomatically colonize the gastrointestinal tracts of humans and animals [1]. Infections are commonly triggered after antibiotic ablation of metabolically competing commensals, allowing the pathogen to rapidly expand [2]. As *C. difficile* populations exceed the carrying capacity of the gut nutrient environment, toxigenic strains produce the enterotoxins A and B, and binary toxin in some epidemic strains, which cause severe watery diarrhea and colitis, and result in additional host-origin nutrients entering the gut lumen [3].

*Clostridioides difficile* infection (CDI) is the most common healthcare-associated infection [4] and causes substantial mortality and morbidity with an estimated 500,000 cases per year and a total CDI-attributable expense of over $5 billion in the United States [5, 6]. Recurrent CDI often follows antibiotic exposure and leads to alternative treatments such as, fecal microbiota transplantations and biotherapeutic agents. Therapies targeting *C. difficile’s* metabolism provide approaches to impede *C. difficile’s* colonization and expansion in the gut, to both aid in preventing primary as well as recurrent infections [7].

*C. difficile* primarily utilizes Stickland fermentation of leucine, glycine, and proline in reductive metabolism to support corresponding high-flux oxidative and energy-generating metabolism of substrates including carbohydrates and other amino acids [8–10]. Proline is a preferred reductive substrate with gut pools that originate from the diet other microobes, and from the host, including via collagen breakdown that releases hydroxyproline [11–14].

The pathogen carries metabolic machinery to convert many substrates to proline [15, 16]. Hydroxyproline, commonly from host collagen, is converted to L-proline by 4-hydroxyproline reductase, pyrroline-5-carboxylatereductase, and racemization by proline racemase to D-proline[17]. Ornithine is also readily converted to proline by ornithine cyclodeaminase [9].

For reductive metabolism *C. difficile’s* proline reductase selenoenzyme reduces D-proline to 5-aminovalerate [18], with coupling to the Rnf complex to generate proton gradients [19]. Proline reductase is encoded by the *prd* operon by seven genes, *prdC*, *prdR*, *prdA*, *prdB*, *prdD*, *prdE,* and *prdF.* The *prdR* transcriptional regulator responds to proline to induce *prd* operon expression [15, 19]. The PrdB catalytic and selenium-containing subunit is essential for enzyme activity [20].

Prior studies of a *ΔprdB* mutant in the CD630 strain identified *in vitro* functions of the gene on proline fermentation [18], Bouillaut *et al*. also demonstrated maintenance of growth *in vitro* by the *prdB* mutant, suggesting compensatory recruitment of other metabolic pathways to overcome the absence of proline reductive metabolism [15]. *In vivo* colonization studies in mice constituted with donor human microbiota indicated defects in early colonization and toxin production [21] [22], though without capacity to assess impact on the development of symptomatic disease as the strain is non-infective in mice. Per prior limitations to genetically manipulate mouse-infective strains, the enzyme’s role in an *in vivo* context of active infection has not been well-defined, including broader effects on pathogen metabolism and recruitment of cellular systems that modulate growth and virulence programs.

We deleted the *prdB* gene in the mouse infective strain ATCC43255 to evaluate its *in vivo* role in the pathogen’s redox metabolism, virulence and host outcomes in highly susceptible gnotobiotic mice. We demonstrate how absence of proline reductase metabolism causes broad disruptions in the pathogen’s ability to efficiently recruit growth-promoting redox metabolism, impeding early events in pathogen colonization and growth, to extend host survival but not prevent the development of severe disease. Our findings identify proline reductase’s contribution to central metabolic strategies used by *C. difficile* in early gut colonization, growth, and eventual virulence, and identify metabolic strategies to aid in preventive and therapeutic strategies.

## Results

Absence of reductive proline metabolism delays *C. difficile* colonization and growth in an enriched gut nutrient environment.

To evaluate the functions of *C. difficile’s* proline reductase metabolism in disease for a susceptible host, 6-week old Swiss-Webster gnotobiotic mice infected with 1000 spores of WT ATCC43255. Mice infected with the wild-type strain rapidly succumbed over 48-72 hours (Figures 1A-B). In contrast, mice infected with the Δ*prdB* mutant showed extended host survival by 24h, with one mouse surviving to 14 days post-infection. Median survival times for the WT and Δ*prdB* groups were 1.85 d and 2.85 d, respectively (Figure 1B).

**Figure 1.**
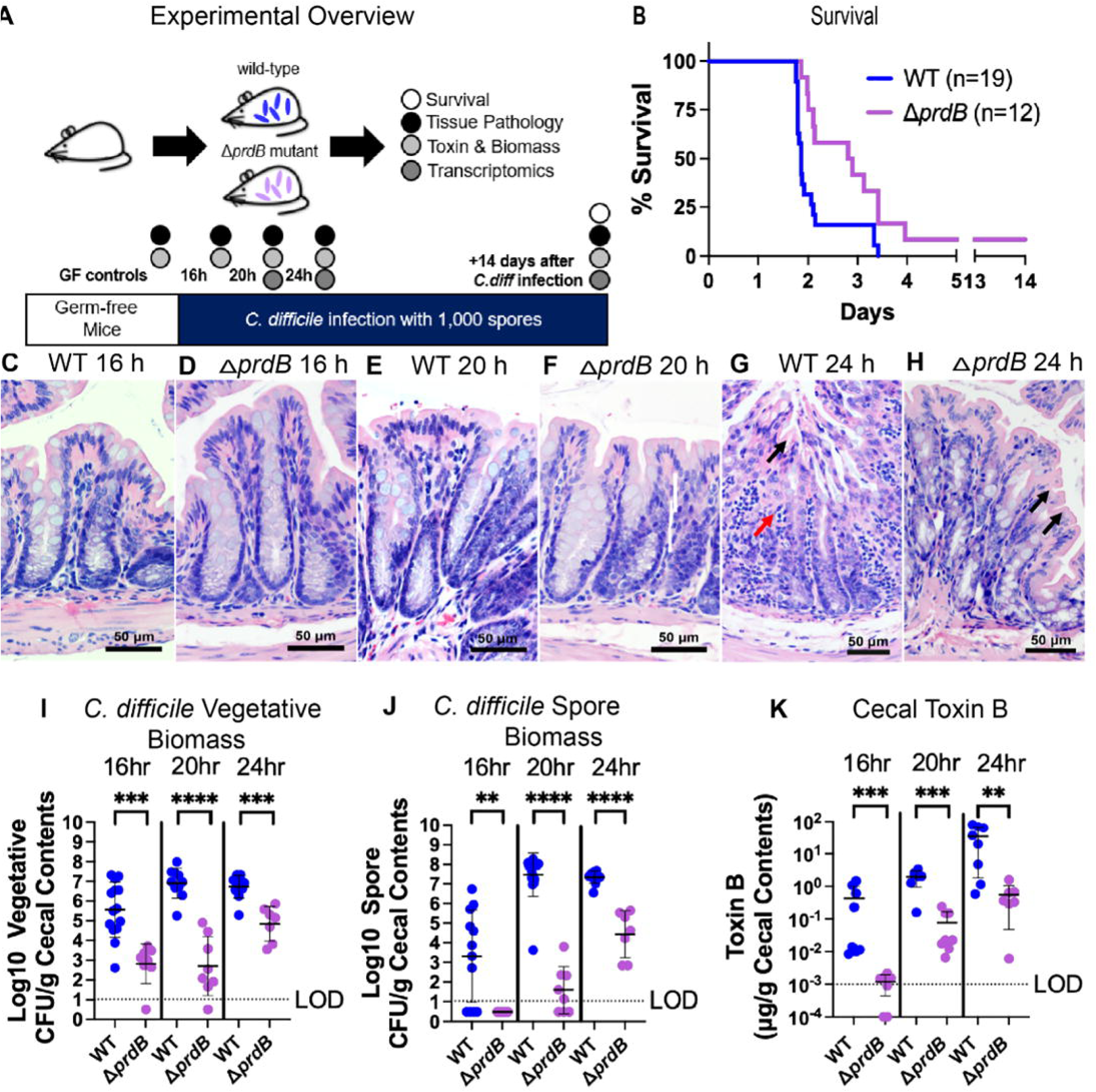
Mouse infection with a *C*. *difficile* Δ*prdB* mutant shows delayed growth and toxin production *in vivo*. **A)** Experimental overview showing the gnotobiotic *C. difficile* infection model. **B**) Kaplan-Meier survival curves showing delayed lethal infection in mice infected with the Δ*prdB* mutant, Mantel-Cox significance p = 0.007. **C-H**) Hematoxylin and eosin (H&E)-stained colon sections. Magnification 200X, scale bar 50 μm. **C**) Colonic mucosa from a germ-free mouse infected with wild-type (WT) *C. difficile* at 16 hr of infection showing no signs of disease, **D**) Healthy colonic mucosa from a germ-free mouse infected with the Δ*prdB* mutant at 16 h of infection, **E**) Colonic mucosa from a germ-free mouse infected with WT *C. difficile* at 20 h of infection**, F**) Colonic mucosa from a germ-free mouse infected with the Δ*prdB* mutant at 20 h of infection **G**) Colonic mucosa from a germ-free mouse infected with WT *C. difficile* at 24hr showing surface epithelial stranding and sloughing (black arrow) and transmural neutrophilic infiltrates (red arrow), and **H**) Colonic mucosa from a germ-free mouse infected with the Δ*prdB* mutant at 24 h of infection showing slight disease and vacuolation of surface colonocytes (black arrows). **I**) Log_10_ *C. difficile* vegetative biomass. **J**) Log_10_ *C. difficile* spore biomass. **K**) Cecal toxin B (μg toxin/g cecal contents). Bars show mean and standard deviation. Limit of detection denoted by LOD. Mann-Whitney significance values: ∗∗ 0.001 < p ≤ 0.01, ∗∗∗ 0.0001 < p ≤ 0.001.

Histologically, by 16h post-challenge the colonic mucosa in the WT-infected mice showed evolving transmural inflammation with neutrophil recruitment into the mucosa (compare Figures 1C-D). In contrast, the colonic mucosa appeared normal at 16h in mice challenged with the *ΔprdB* mutant (Figure 1E). However, over 20-24h of infection, vacuolation and stranding of surface colonocytes occurred with the *ΔprdB* mutant, while more severe surface epithelial sloughing and transmural neutrophilic infiltrates were visible by 24 h post-infection with the WT strain (Figure 1G).

Assessments of cecal toxin B, and *C. difficile* vegetative and spore biomass demonstrated a colonization defect in the *ΔprdB* mutant with decreased spore and vegetative biomass, and delayed production of toxin B in cecal contents throughout infection (Figure 1I-K).

Ablation of reductive proline metabolism broadly disrupts *C. difficile’s* metabolic strategies to colonize and adapt to changing gut nutrient conditions.

*In vivo* analyses of *C. difficile* gene expression in cecal contents identified multiple metabolic disruptions over the course of infection with broader impact on additional cellular systems (Figure 2). Over 16-20h of colonization, in contrast to the wild-type strain, the *ΔprdB* mutant failed to induce its Stickland oxidative fermentation and ornithine fermentations pathways that generate alanine and proline (Figure 3). To compensate, the mutant induced expression of reductive Stickland leucine and glycine metabolism (Figure 2), in addition to the other *prd* operon genes, though with undetectable *prdB* (Figure 3). The mutant compensated for the defects in its Stickland redox metabolism by recruiting pathways for sugar alcohol metabolism, inducing mannitol transport and fermentation at 20h, followed by sorbitol metabolism at 24h post-infection, respectively (Figure 3). Fructose transport and fermentation genes, *fruABC* and *fruK,* were also significantly induced in *prdB* mutant, as compared to the wild-type strain, at 20h post-infection (Figure 3).

**Figure 2.**
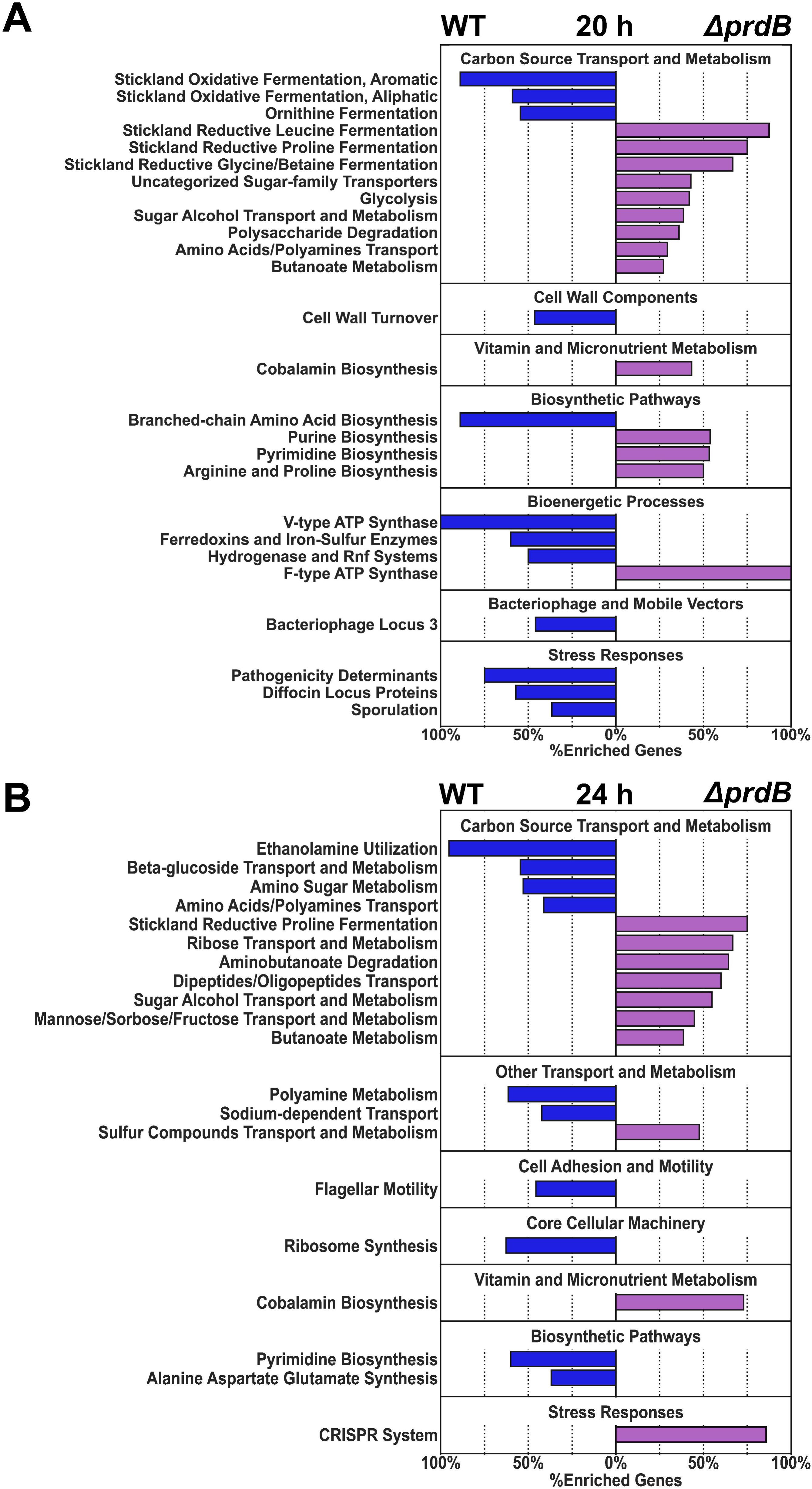
*C*. *difficile* Δ*prdB* mutant induces metabolic disruptions *in vivo* reflected by the pathogens global gene expression. Metatranscriptomic analyses of *C. difficile’s* metabolic, core cellular machinery and cellular stress systems that were significantly enriched in mice challenged with wild-type *C*.*difficile* (WT, blue) or Δ*prdB* mutant *C*. *difficile* (Δ*prdB,* purple). Horizontal bars indicate the percentage of enriched genes with a Benjamini-Hochberg-corrected p-value ≤ 0.05. Headings above the bars show general pathway categories with subpathways on the left. (A) Enriched pathways at 20 h post-infection with WT or Δ*prdB C*.*difficile.* B) Enriched pathways at 24 h post-infection with WT or Δ*prdB C*.*difficile*.

**Figure 3.**
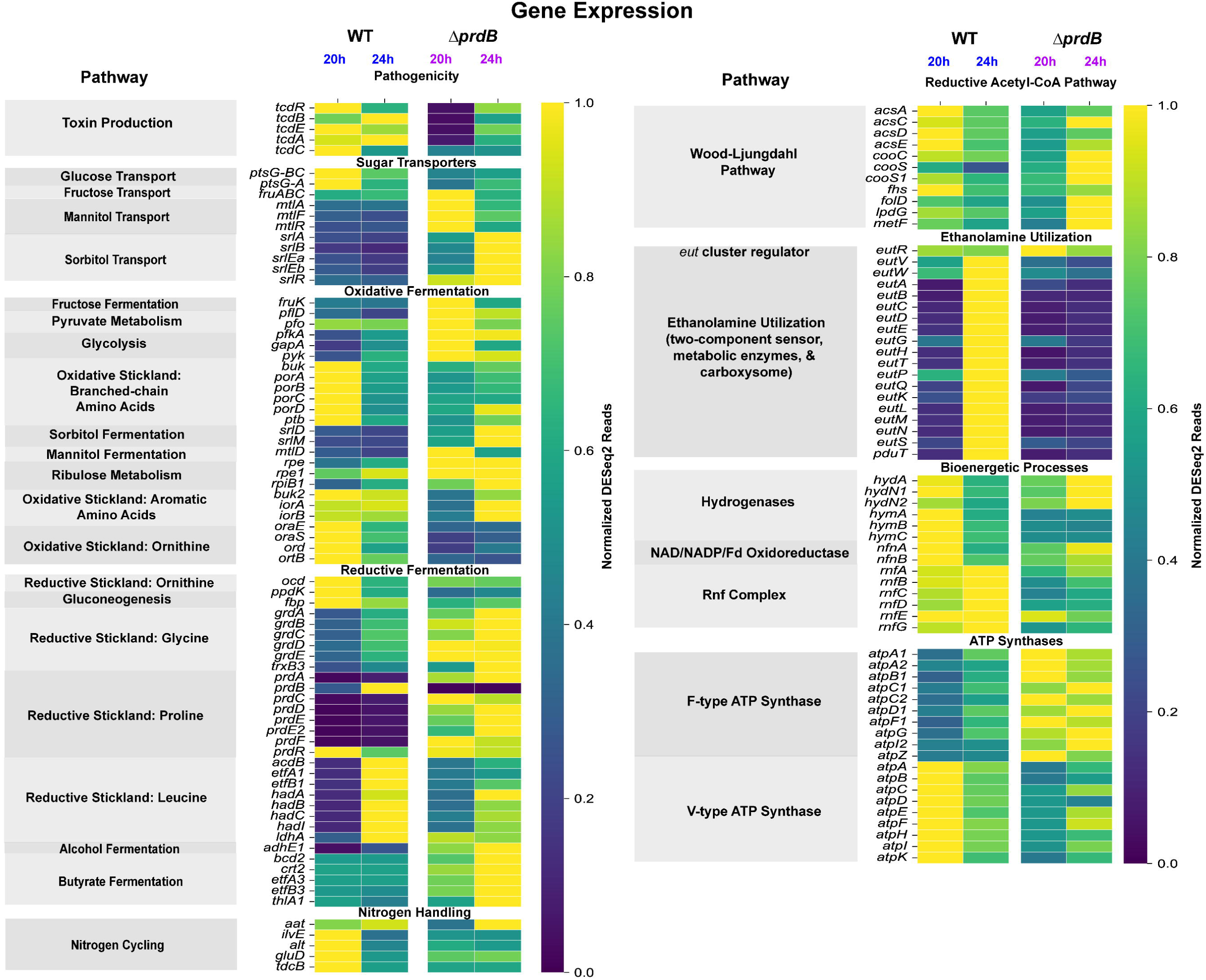
*C*. *difficile* Δ*prdB* mutant fails to induce energy-generating oxidative Stickland metabolism and adjusts to alternative reductive amino acid and oxidative carbohydrate fermentation systems. Heatmap colors indicates the proportion of maximum gene expression for each gene as normalized DESeq2 reads (scale 0-1). Timepoints are indicated at 20-and 24 h post-infection with either the *C*. *difficile* wild-type (WT, blue) ATCC43255 strain or Δ*prdB* mutant (Δ*prdB,* purple). Pathogenicity and metabolic categories (*e.g*., oxidative fermentation) are listed between each break in the heatmap and pathways are labelled in the gray boxes to the left of the gene names.

As toxin-mediated damage to mucosal surfaces evolved in mice, the WT strain induced ethanolamine fermentation by 24h of infection [23], a fermentable substrate released from damaged mucosa and from host membrane-associated phosphatidyl-ethanolamine lipids (Figure 2B). In contrast, the Δ*prdB* mutant consistently relied on glycolytic and mixed acid fermentation reactions that produce acetate and butyrate, particularly over 20-24h of infection, and showed limited metabolic adaptations, inducing only pathways for the transport and metabolism of ribose, transformation of mannose for glycolysis, oxidative Stickland pathways for aromatic amino acids, and upregulation of pathways for cobalamin biosynthesis (Figure 2B) [24]. With the underlying defects in Stickland metabolism, the *prdB* mutant showed reduced expression of its alanine transaminase that integrates glycolytic and de-aminating Stickland metabolism to aid in the generation of anabolic substrates for protein and cell wall synthesis (Figure 3) [25]. Systems associated with bioenergetic processes, including hydrogenases, V-type ATP synthases, and Rnf complex were also repressed in the mutant (Figure 3). The mutant strain also demonstrated significantly reduced expression of sporulation, ethanolamine utilization, and flagellar genes at 24h of infection.

The combined *in vivo* metatranscriptomics, with host and *C. difficile* phenotypes, illustrated how sole disruption of the *ΔprdB* gene and reductive proline metabolism broadly impacted the pathogen’s early recruitment of efficient redox metabolism and subsequent phenotypes on delayed growth, sporulation, and toxin production. Despite these delays in growth and virulence, the pathogen eventually produced sufficient toxin to cause severe disease in monocolonized mice.

## Discussion

We generated a Δ*prdB C. difficile* mutant to define the *in vivo* functions of reductive proline metabolism in the pathogen’s broader redox metabolism, growth, and virulence. The Δ*prdB* mutant demonstrated delayed colonization, growth, and toxin production which extended host survival [26]. While gnotobiotic mice monocolonized with the mutant ultimately succumbed to infection, our findings demonstrated how the absence of reductive proline metabolism impacted *C. difficile’s* capacity to efficiently recruit and harness its redox metabolism for energy and anabolic substrates during early stages of gut colonization. By 24h of infection, Δ*prdB* mutant expressed comparable levels of its Pathogenicity Locus genes *tcdA*, *tcdB*, *tcdE,* and *tcdR* to wild-type, explaining the delay in symptoms and survival, with the resulting host outcome of severe disease. However, even by 24h, as host symptoms progressed, the Δ*prdB* mutant still failed to adequately harness available gut nutrients in gnotobiotic mice.

With the disruption of proline reductase metabolism, the pathogen recruited alternative Stickland reductive pathways for leucine and glycine fermentation [15]. Notably, the mutant failed to induce ornithine metabolism genes during early stages of colonization including those converting ornithine to alanine, a key growth-promoting substrate, as well to proline by ornithine cyclodeaminase [27]. While the *prdB* mutant recruited pathways for glycolysis, pyruvate metabolism, and subsequent butyrate metabolism, it did not concomitantly induce its alanine transaminase to support efficient amino group nitrogen handling from deaminating Stickland metabolism, including for reductive glycine and leucine metabolism.

Absence of reductive proline metabolism shifted *C. difficile* into become a more glycolytic pathogen, but one lacking optimized redox metabolism to efficiently grow on carbohydrate substrates. Lacking efficient oxidative and reductive Stickland metabolism during early stages of colonization, the Δ*prdB* mutant turned to alternative carbohydrate sources including sugar alcohols, fructose, mannose, and ribose. While mannitol originates from dietary sources [9, 28, 29], sorbitol can derive from dietary and host sources, including from immune cells via the activity of aldose reductase, an enzyme that is upregulated during host inflammation [30]. The mutant also induced multiple cobalamin biosynthetic operon genes to support cobamide-dependent pathways [31].

Our findings define complex mechanisms by which reductive proline metabolism enables *C. difficile’s* initial colonization, growth, and toxin production by supporting efficient induction of broader energy-generating and anabolic pathways. While the *ΔprdB* mutant’s metabolic shifts to glycolytic and mixed acid fermentations supported eventual growth and pathogenesis in a highly enriched gut nutrient environment, its remained unable to efficiently harness preferred amino acids and newly available nutrients, such as ethanolamine for energy and growth as infection progressed. Given the presence of proline reductase metabolism in other *Clostridial* pathogens, such as *C. botulinum,* strategies to disrupt this pathway provide opportunities to leverage metabolic approaches in treatment, and for disease prevention.

## Materials and Methods

### Construction of the *ΔprdB* mutant in ATCC43255

Gene deletion in *C*. *difficile* was performed as described by Peltier et al. (2020) [32]. Regions upstream (F-TTTTTTGTTACCCTAAGTTTGGTACATTAGGAGCTCAAC, R-AAACGTGAGCCCTATCCATTATAATTCTACCTC) and downstream (F-AATGGATAGGGCTCACGTTTAAAATGTAATTTATATTTG, R-AGATTATCAAAAAGGAGTTTGGAGATATAATCATAGGTACATTTC) of the genes of interest were PCR-amplified with subsequent PCR to verify the deletion of *prdB* in *C. difficile* ATCC 43255 (F-ATCC43255 GAAATAATGGGACAAGGTG, R-ATCC43255

CCTTTGTTCTTAAAACCCC). PCR fragments and linearized pDIA6754 were mixed and assembled using Gibson Assembly (NEB, France) and transformed by heat shock in *E. coli* NEB 10ꞵ strain [32]. The plasmid constructions were verified by sequencing and plasmids with the right sequences were transformed in *E. coli* HB101 (RP4). The resulting strains were used as donors in a conjugation assay with the relevant C. difficile strains. Deletion mutants were then obtained using a counter-selection [32].

### *In vivo* infection studies in gnotobiotic mice

All gnotobiotic mouse studies in the Massachusetts Host-Microbiome Center at BWH were conducted under IACUC protocol 2016N000141. Equal numbers of male and female Swiss webster gnotobiotic mice at 6-7 weeks old were singly housed in sterile OptiMice cages (Animal Care Systems, Centennial, CO). Germ-free mice were challenged with either 1 x 10^3^ spores of wild-type ATCC 43255 or Δ*prdB* mutant. Mouse survival up to 14 d post-infection with *C. difficile* was monitored for disease progression. Body condition scoring was utilized to assess nutritional status and disease progression [33]. Death was not used as an endpoint for disease.

In timepoint studies at 16, 20 and 24h post-challenge, cecal contents were immediately collected and frozen in liquid nitrogen for downstream biomass counts, transcriptomics, and toxin B ELISA analyses, while the gastrointestinal tract was fixed in formalin for paraffin embedding and hematoxylin and eosin (H&E) staining of 5mm sections.

### Quantification of *C. difficile* biomass

Previously flash-frozen cecal contents were weighed in 1.5 ml tubes and kept frozen on a cold block prior to transfer into a Coy anaerobic chamber (Coy Labs, Grass Lake, MI) at 37°C with an atmosphere of 2.5% H_2_, 10% CO_2_, 87.5% N_2_. In the chamber, samples were homogenized in 1 mL of pre-reduced PBS with 40 mM cysteine as a reducing agent (Millipore-Sigma, St. Louis, MO), serially diluted, and plated in triplicate to *C. difficile*-selective CHROMID® agar (Biomérieux, Durham, NC). *C. difficile* colonies were identified as large black colonies and counted at 72 h post-incubation.

*C. difficile* spores were isolated from pre-weighed, diluted, and homogenized cecal contents by exposure to 50% ethanol for 1 hr and subsequently centrifuged to form a pellet. The pellet was washed twice with PBS to remove the ethanol prior to serial diluting and plating the samples in triplicate to CHROMID® agar. *C. difficile* colonies from the spore fraction were identified and counted as described above.

### Histopathology

Formalin-fixed colon tissues from germ-free, *C. difficile* mono-colonized animals were blocked, paraffin-embedded, and cut into 5 mm sections for staining with hematoxylin and eosin (H&E) stain (H&E; Thermo-Fisher, Waltham, MA) as previously described [34]. Slides were visualized using a Nikon Eclipse E600 microscope (Nikon, Melville, NY) to assess features of epithelial damage including cellular stranding, vacuolation, mucosal erosions and hyperplasia as previously described [26].

### Toxin B ELISA

Toxin B concentrations from cecal contents were measured as previously described by Zarandi et al. 2017 and Girinathan et al., 2021 with the following adjustments [26, 35]. Microtiter plates were coated with 5 µg/ml of the capture antibody (MAB13508, Abnova, Heidelberg, Germany). Plates were washed three times with PBS-Tween20 and blocked at RT for 2 h with SuperBlock (Thermo Scientific, Waltham, MA). Cecal contents were diluted with 1X PBS, vortexed, and centrifuged at 17,000 x g for 1 min. The sample supernatants and toxoid B standard controls (32-0 ng/ml) (BMLG1550050, Enzo Life Sciences, Southold, NY) were tested in triplicate. Plates were incubated with a biotinylated detection antibody (ab252712, Abcam, Cambridge, UK) and Pierce High-Sensitivity Streptavidin-HRP (Thermo Scientific, Waltham,MA). Plates were washed with PBS-Tween20 between incubations. TMB substrate (Thermo-Fisher, Waltham, MA) was used for signal detection at 450 nm with a BioTek Synergy H1 plate reader (Biotek Instruments Inc., Winooski, VT). Values were interpolated in GraphPad Prism 9 (GraphPad, San Diego, CA) to calculate toxin B concentrations. Significant differences between treatment groups were evaluated by non-parametric Mann-Whitney test with differences declared at a p-value ≤0.05.

### *In vivo* Bacterial RNA Sequencing

Snap-frozen cecal contents were weighed into lysis tubes with 0.1 and 0.5 mm beads (Zymo Research, Irvine, CA) and 750 µl DNA/RNA Shield to inactivate nucleases. Cells were lysed at 4 °C for a total of 30 min. on a Vortex Genie2 (Scientific Industries, Bohemia, NY) fitted with a 2 ml tube holder (Qiagen, Germantown, MD). RNA was purified with the ZymoBIOMICS RNA Miniprep kit (R2001, Zymo Research, Irvine, CA). Purified RNA was concentrated with the RNA Clean and Concentrator-5 kit (R1013, Zymo Research, Irvine, CA) with a second DNAse I step. The concentration and integrity of the purified RNA were quantified with the Qubit 3 fluorometer (Thermo Scientific, Waltham, MA) and 2100 Bioanalyzer (Agilent Technologies, Lexington, MA), respectively. Prior to library prep, qPCR was employed to assess DNA contamination with DNA polymerase III-specific primers (F: 5’-CCCAACTCTTCGCTAAGCAC-3’; R: 5’-TCCATCTATTGCAGGGTGGT-3’). The Zymo-Seq RiboFree Total RNA Library Kit (R3003; Zymo Research, Irvine, CA) was used to generate cDNA libraries following the manufacturer’s instructions. The molar concentration of each individual cDNA library was calculated from the Qubit concentration (ng/µl) and bioanalyzer length. Each sample was diluted to 4 nM and equimolar pooled. Sequencing on the Illumina MiSeq with the MiSeq 150-cycle v3 kit (MS-102-3001; Illumina, San Diego, CA) was used to assess bacterial versus host enrichment of the libraries. Bacterial read proportions calculated from Bowtie 2 and HTSeq analyses were used to estimate individual sample inputs required to obtain a minimum of 1 million reads for each bacterial species within a sample [26]. A second 4 nM pool was generated and sequenced on the Illumina NovaSeq 6000 with the NovaSeq 6000 S4 Reagent v1.5 (300 cycles) kit (20028312, Illumina, San Diego, CA) at the Molecular Biology Core Facilities at the Dana-Farber Cancer Institute (Boston, MA). Data processing and enrichment analyses of the transcriptomes was performed as described by Girinathan et al. 2021[26].

